# Chromatiblock: scalable whole-genome visualization of structural differences in prokaryotes

**DOI:** 10.1101/800920

**Authors:** Mitchell John Sullivan, Harm van Bakel

**Affiliations:** Department of Genetics and Genomic Sciences, Icahn Institute for Data Science and Genomic Technology, Icahn School of Medicine at Mount Sinai, New York, NY 10029, United States of America

## Abstract

**Summary:** Chromatiblock is a Python application for visualizing the presence, absence and arrangement of syntenic blocks across large numbers of complete bacterial genomes.

**Availability:** Chromatiblock is freely available under a GPL license, for macOS, GNU/Linux and and Microsoft Windows from https://github.com/mjsull/chromatiblock/

## 1 Introduction

Visualizing structural variation between complete prokaryotic genomes is important for identifying the genetic basis of strain differences. This is generally accomplished by displaying the results of serial pairwise comparisons or multiple alignments in linear or circular layouts. Serial pair-wise comparisons can be created using tools such as Easyfig (Sullivan, et al., 2011) or GenoplotR (Guy, et al., 2010) that display linear pairwise comparisons between two or more genomes. However, genomic loss, gain and structural variation can only be directly inferred for adjacent genomes. Multiple alignment visualization tools such as Mauve (Darling, et al., 2004) solve this issue by representing syntenic regions as linear blocks and using lines to connect blocks across genomes. In large figures this can result in crisscrossing lines that are often difficult to interpret. Alternatively, ring plots, such as those created by the BLAST ring image generator (BRIG) (Alikhan, et al., 2011) or the CGView Comparison Tool (CCT) (Grant, et al., 2012) use a series of concentric circles to display the presence or absence of genomic regions across multiple genomes. These regions are ordered according to a reference, and as such they convey no information about their arrangement in each non-reference genome. Alternatively, Circos (Krzywinski, et al., 2009) plots show genomes around the outside edge of a circle and represents regions of similarity as arcs, but this approach scales poorly as the number of arcs increases exponentially with each genome. Representing many genomes as circles can also result in large size differences between inner and outer rings, further complicating interpretation.

Here we present Chomatiblock, an application for visualizing syntenic blocks in multiple genome alignments. Chromatiblock was designed to create a linear visual representation of structural variation, including the presence and absence of genomic regions in an easy-to-comprehend and scalable manner, adding to the visualization options available for alignments of large numbers of complete genomes.

## 2 Implementation

Chromatiblock is a Python script available under a GPL license and runs on macOS, GNU/Linux and Microsoft Windows operating systems. Chromatiblock can be used to create publication-quality images displaying arrangement and presence of syntenic blocks. The results can also be viewed as an interactive webpage that allows the user to zoom, pan and highlight shared regions across genomes.

Chromatiblock takes an extended multi-fasta alignment (MAF) files as input, which can be generated by a variety of multi-genome alignment programs (Angiuoli and Salzberg, 2011; Minkin and Medvedev, 2019). Alternatively, when provided with FASTA-formatted files for a set of genomes of interest, Chromatiblock can run Sibelia (Minkin, et al., 2013) to automatically generate the required input. Once syntenic blocks have been identified in the MAF file, Chromatiblock will generate a dual-panel layout consisting of a global alignment view and a detailed view of regions that differ between genomes. The global alignment view shows the arrangement of core blocks (i.e. syntenic regions found once in all genomes) in the alignment and how non-core blocks (i.e found in 2 or more genomes) and unique sequences (i.e. found in a single genome) are arranged relative to the core blocks. Core blocks are aligned according to their arrangement in the first genome. The color the core blocks for each genome is determined by its position. Between each core block there exists a combination of non-core blocks and unique sequence. This combination is grouped and positioned between the two core blocks to which they are adjacent. In instances where the group cannot be placed between its two adjacent core blocks it is placed arbitrarily next to one of the core blocks to which it is adjacent. This is indicated by removing the gap between core and non-core blocks. An example of a global alignment of 28 complete *C. difficile* genomes is shown in **Fig. 1A**. A large inversion can be observed in the third isolate from the top, indicated by a difference in ordering of core block colors relative to the reference. Plasmids, found in 9 genomes, consist entirely of non-core and unique blocks. They are positioned on the right side of the figure. The presence or absence of specific user-provided gene sequences can also be indicated by distinct gene symbols and are automatically annotated using BLASTx. In the example, six isolates contain a transposon carrying the *erm(B)* gene, encoding a 23S rRNA methyltransferase that confers resistance to erythromycin. The *erm(B)*gene is also present in an ST54 isolate but located on a novel transposon and inserted elsewhere in the genome (**Fig. 1A**).

**Fig. 1.**
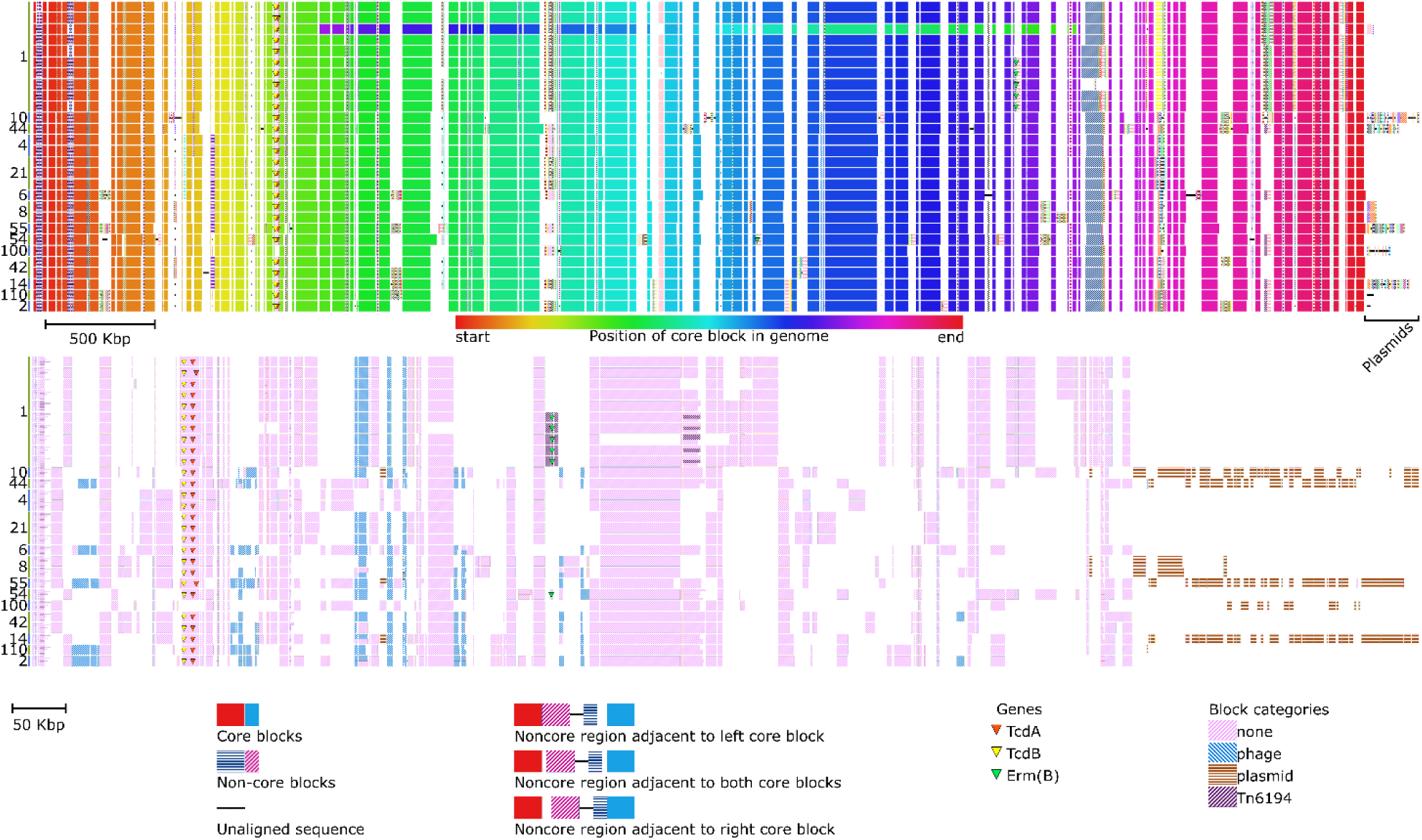
Chromatiblock visualization of 28 *Clostridium difficile* genomes. MLST of each isolate is indicated on the left. **A**) Global alignment view. Core blocks across genomes (rows) are visualized as vertically aligned solid rectangles that are colored according to their position in the genome. Non-core blocksare visualized as patterned rectangles, with each block represented by a unique combination of pattern and color. Finally, sequences unique to a single genome are depicte d as solid black lines.**B)** Alignment difference view. Each genome is represented as a row and each non-core block is assigned a column in the order they are most commonly found in the genome. Presence of each non-core block is shown as a patterned rectangle in the genomes row. As non-core blocks may be present more than once, duplicates are shown by splitting the blocks according to repeat number.

The alignment difference view shows the presence and absence of all non-core blocks. Chromatiblock can use BLAST+ to categorize and color each non-core block based on a user-provided reference database of nucleotide or amino acid FASTA files. Categories can also be assigned based on the size of the contig in which the non-core block is found. The example in **Fig 1B** shows that the main *C. difficile* pathogenicity locus (PaLoc) that contains the genes encoding the TcdA enterotoxin and TcdB cytotoxin, has been lost in the ST100 isolate. Plasmids carried by *C. difficile* are very chimeric, with large regions being shared, but with only the two MLST8 isolates carrying identical plasmids.

In conclusion, Chromatiblock allows users to quickly and easily create publication-quality figures showing structural changes and genetic diversity at the whole genome level.

